# A *Staphylococcus* phage recombination cassette that enables recombineering across distant bacteria

**DOI:** 10.64898/2026.10.30.761037

**Authors:** Pascal Maguin, Maedeh Aghahosseini, Maj Brodmann, Victor Chen, Jeff Zhu, Karolina Szenkiel, Luciano A. Marraffini, Andrew Varble

## Abstract

Modifying bacterial genomes with phage recombinases, a process termed recombineering, has enabled precise genome editing, but its reach is limited by host specificity: each new organism generally requires its own recombinase to be identified and optimized. Using known phage recombination genes as synteny anchors, we mined *Staphylococcus* genomes and recovered a diverse collection of phage recombination cassettes, spanning the RAD52-like annealase superfamily (Sak, Erf) and the RAD51/RecA-like Sak4 family. Functional testing of representatives in *S. aureus* identified several highly efficient systems that support recombineering. We engineered the most efficient of these into a single-plasmid tool that yields markerless editing in *S. aureus* in four days. Notably, where other recombineering systems rely on host-specific factors for nuclease evasion and ssDNA loading, this cassette encodes both activities itself: a predicted broad-spectrum nuclease inhibitor and a dedicated single-stranded DNA-binding protein. We therefore tested the cassette across phylogenetically distant hosts, and observed recombination in *Streptococcus pyogenes, Enterococcus faecium, Francisella novicida*, and *Escherichia coli*. These results identify a self-sufficient class of phage recombination systems and provide a portable tool for genetic manipulation in bacteria that lack established recombineering methods.

## Introduction

Bacteriophages encode their own DNA recombination machinery, and these systems have been repurposed to engineer bacterial genomes in an approach termed recombineering. In recombineering, a linear DNA template is introduced into a cell expressing phage recombination proteins, which shield it from host nucleases and facilitate recombination into the chromosome (1,2). The lambda Red system of *E. coli* is the best-characterized example, and the ease of editing it affords has enabled gene knockout libraries (3), directed protein evolution in bacteria (4,5), and the recoding of entire genomes (6–8). The lambda Red system requires three activities. Exo is a 5′→3′ exonuclease that acts on the transformed linear DNA to generate long 3′ single-stranded overhangs necessary for recombination. Gam binds the *E. coli* RecBCD recombinase complex to shield the transformed linear DNA from degradation by exonucleases (9). The recombinase acts at the replication fork, annealing incoming DNA to the lagging-strand template (10) through a specific interaction with the host single-stranded DNA-binding protein (SSB) that binds the ssDNA generated by Exo (11). Due to the requirement for specific protein-protein interactions between Gam, SSB, and RecBCD, the lambda Red system is not efficient in hosts other than *E. coli*. More generally, portability of recombineering systems in bacterial species depends on compatibility between the phage and host proteins, necessitating the development of individualized systems for each organism (12).

This limitation is evidenced in the lack of tools for the genetic manipulation of bacterial pathogens. *Staphylococcus aureus* is a leading cause of human infection and has acquired resistance to every antibiotic deployed against it (13), yet its genome is still modified largely by plasmid-based allelic exchange, which requires cloning, multiple passages, and the screening of many clones for each edit (14). Oligonucleotide recombineering with a heterologous recombinase works in clinical isolates and can be coupled to Cas9 counterselection (15), and CRISPR editing is efficient once restriction– modification barriers are removed (16). Because direct CRISPR targeting is lethal in bacteria (17,18), however, these approaches select for rare recombinants rather than promote mutagenesis and still depend on the availability of an efficient recombination system (19,20). Edits are consequently made in a small number of laboratory strains and rarely in the clinical isolates that carry the pathogenic repertoire of interest (21–23), a pattern repeated across other bacterial pathogens (24–27).

The success of the lambda Red system has motivated the search for analogous systems that function in organisms where lambda Red is ineffective or that offer greater versatility. These efforts have focused largely on the phage-encoded recombinase itself. Developing new systems often involves identifying a phage recombination cassette from a phage that infects the target organism, often a RecE/RecT homolog (27–31). Because the RecET pair lacks a Gam-like inhibitor of host exonucleases, the systems are restricted to single-stranded oligonucleotide templates that resist degradation upon transformation and rarely approach the efficiency of lambda Red. To identify high-efficiency or broad-spectrum recombinases, large libraries of single-strand annealing proteins (SSAPs) have been screened for their ability to mediate genome editing (32,33), but their use remains dependent on the host’s own nuclease protection and DNA-loading machinery.

Multiple lines of evidence indicate that annealases rarely function as self-contained units when moved between organisms. For double-stranded substrates, RecET and Redα/Redβ each require their cognate exonuclease, and the two pairs cannot be mixed (34). Host compatibility is also limiting where SSAPs recognize the C-terminal tail of the host SSB and are portable only where that interaction is preserved (11). Consistent with this, the annealing activity of Sak4 is strongly stimulated by its cognate phage SSB through the SSB C-terminus (35). Furthermore, the BAS operon of *P. aeruginosa* phage Ab31, which natively encodes an annealase, exonuclease, and SSB, required addition of an exogenous RecBCD inhibitor for maximal recombineering in *Pseudomonas*, while its SSB independently enhanced recombination (36). Contiguous annealase– exonuclease pairs have been mined and tested across genera (12,37–39), and some operons carry a putative exonuclease inhibitor as a third gene (24), but no systematic search has treated a nuclease inhibitor, recombinase, and cognate SSB as a single transferable cassette.

We reasoned that these constraints could be overcome by a recombination module that supplies both activities, namely a nuclease inhibitor with broad specificity rather than a single protein-protein partner, together with its own dedicated SSB. Such a self-sufficient cassette would act independently of host factors and edit double-stranded DNA in phylogenetically distant bacteria. To find one, we used known phage recombination genes as synteny anchors to mine *Staphylococcus* genomes, recovering a diverse collection of phage recombination cassettes that span distinct recombinases of the RAD51 (Sak4) (35) and RAD52 (Sak, Erf) single-strand annealing protein superfamilies (40,41). Functional testing of representatives in *S. aureus* identified several systems that support efficient recombineering, and we engineered the most active into a single-plasmid tool that generates nearly markerless gene deletions in four days. Notably, because this cassette encodes a broad-spectrum nuclease inhibitor and a dedicated SSB that do not require additional host factors to function, it supports recombination not only in *S. aureus* but also in *Streptococcus pyogenes, Enterococcus faecium, Francisella novicida*, and *Escherichia coli*. These results define a self-sufficient class of phage recombination systems and provide a portable platform for genome editing in bacteria that lack established recombineering methods.

### SapSak4 recombineering

Most *S. aureus* phages encode a two-gene recombination operon comprising an SSAP/recombinase and a cognate SSB (42). Both proteins are required for phage replication in all tested phages (42). To determine whether these operons can mediate recombineering, we sought to isolate the recombination cassette from the prototypical *S. aureus* phage □11 (Figure 1A), a 45-kb prophage carried in strain NCTC 8325 (43). 11 encodes an SSAP of the Sak4 family, which we designate SapSak4 (*S. aureus* prophage Sak4), together with an SSB. Immediately upstream of SapSak4 is a third unannotated gene, which aligns with a Mu-like Gam protein family (Figure 1B). Mu Gam was originally named for its ability to complement deletions in the *gam* gene of lambda phage (44), an essential component of the lambda Red recombineering system. Despite this functional complementation, Mu Gam obstructs host DNA exonucleases by binding and protecting free dsDNA ends (45–48), in contrast to lambda Gam’s direct inhibition of RecBCD (9). Together, these observations indicate that the complete recombination cassette from phage □11 comprises a Mu Gam-like protein to protect dsDNA from host nucleases and SapSak4 together with its cognate SSB to mediate recombination (hereafter, the SapSak4 recombination cassette).

**Figure 1.**
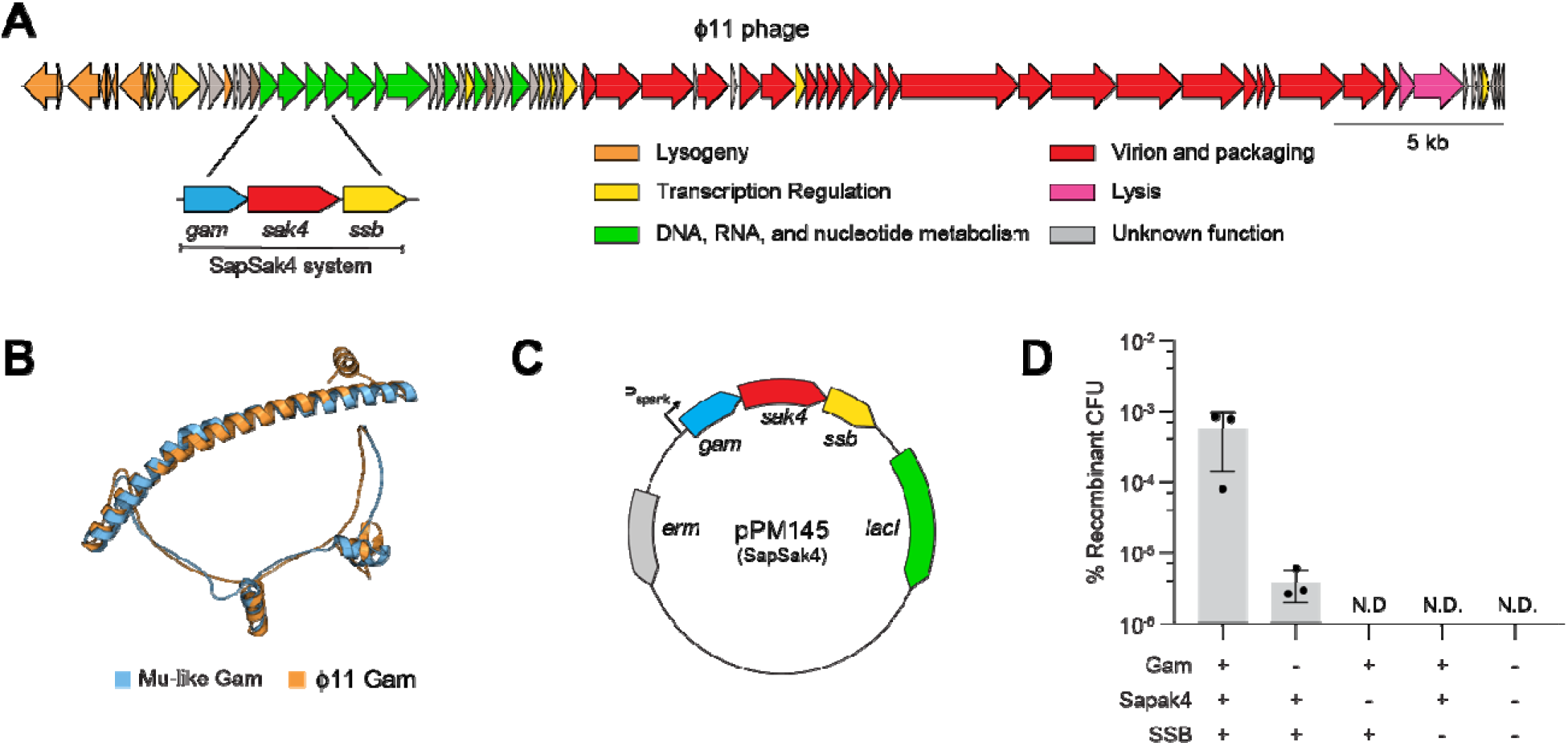
The SapSak4 cassette mediates recombineering in S. *aureus* RN4220. **(A)** Schematic of the□ 11 genome highlighting the location of *gam*, *sak4*, and *ssb*. **(B)** Superimposition of the Mu-like Gam structure 2P2U (blue) and the predicted structure of□11’s Mu-like Gam (orange). **(C)** Schematic of recombineering vector pPM145. The plasmid contains an IPTG-inducible operon with□11’s *gam*, *sak4*, and *ssb*. **(D)** Percent of recombinant CFU following electroporation of a dsDNA substrate containing an antibiotic-resistance gene into *S. aureus* RN4220 harboring a vector with different *gam*, *sak4*, and *ssb* gene combinations or an empty plasmid control. Means of three biological replicates ± SD are reported.

To test whether this three-gene cassette could mediate recombination of dsDNA, the operon was cloned into an inducible *S. aureus* expression vector (Figure 1C). We also constructed a dsDNA recombination template carrying an antibiotic-resistance cassette flanked by 500 bp of homology to the *sau1hsdR* locus of the *S. aureus* chromosome (Supplementary Figure S1A). Electroporating this template into cells carrying the three-gene operon yielded antibiotic-resistant colonies (Figure 1D), and PCR across the expected insertion site confirmed that editing had occurred (Supplementary Figure S1B). Deletion of either *sak4* or the *ssb* from the construct abolished editing entirely, while deletion of *gam* reduced recombineering efficiency by two orders of magnitude (Figure 1D). SapSak4 and the SSB are therefore essential for recombineering, with Gam contributing substantially to efficiency. When the plasmid was induced in the absence of antibiotic selection, it was lost from all edited colonies (Supplementary Figure S1C), possibly reflecting toxicity of the induced system. However, this plasmid loss provides a convenient route to plasmid curing after editing without the need for a secondary counterselection mechanism. Together, these results establish the SapSak4 recombination cassette as a functional recombineering system in *S. aureus*. The genetic requirements suggest a division of labor: SapSak4 and its cognate SSB mediate annealing of the incoming DNA, while Gam improves efficiency without being strictly required, as expected for its predicted nuclease-protection function.

### Mining *Staphylococcus* genomes for phage recombination cassettes

Phage genomes are functionally modular: genes for replication, structure, and lysis are organized into discrete clusters in which gene order is often preserved (49–51). Recombination cassettes follow this pattern, with the Gam–SSAP/recombinase–SSB arrangement commonly observed across Gram-positive phages (52). We therefore exploited this synteny to identify recombination cassettes in staphylococcal phages, and to compare them with the SapSak4 system from□11.

To survey these systems across *Staphylococcus*, we developed an iterative, synteny-guided search strategy that uses individual cassette genes to identify neighboring locus components (Figure 2A). Beginning with the □11 Gam protein sequence, we identified its homologs across *Staphylococcus* genomes and, for every hit, retrieved the immediately flanking genes to capture the associated SSAP/recombinase and SSB. The major SSAPs/recombinases recovered from this round were then used as queries for additional Gam proteins, which in turn seeded searches for further SSAPs/recombinases. Alternating between the *gam* and *SSAP/recombinase* positions allowed the search to span protein families that share no detectable sequence similarity and was repeated until no new SSAP/recombinase or Gam families emerged. Among loci containing potential Gam homologs, 86% also carried an adjacent SSAP/recombinase, and our search recovered 5,603 complete loci containing Gam, a SSAP/recombinase, and an SSB, confirming that the Gam–SSAP/recombinase–SSB cassette is a tightly conserved syntenic unit across staphylococcal prophages.

**Figure 2.**
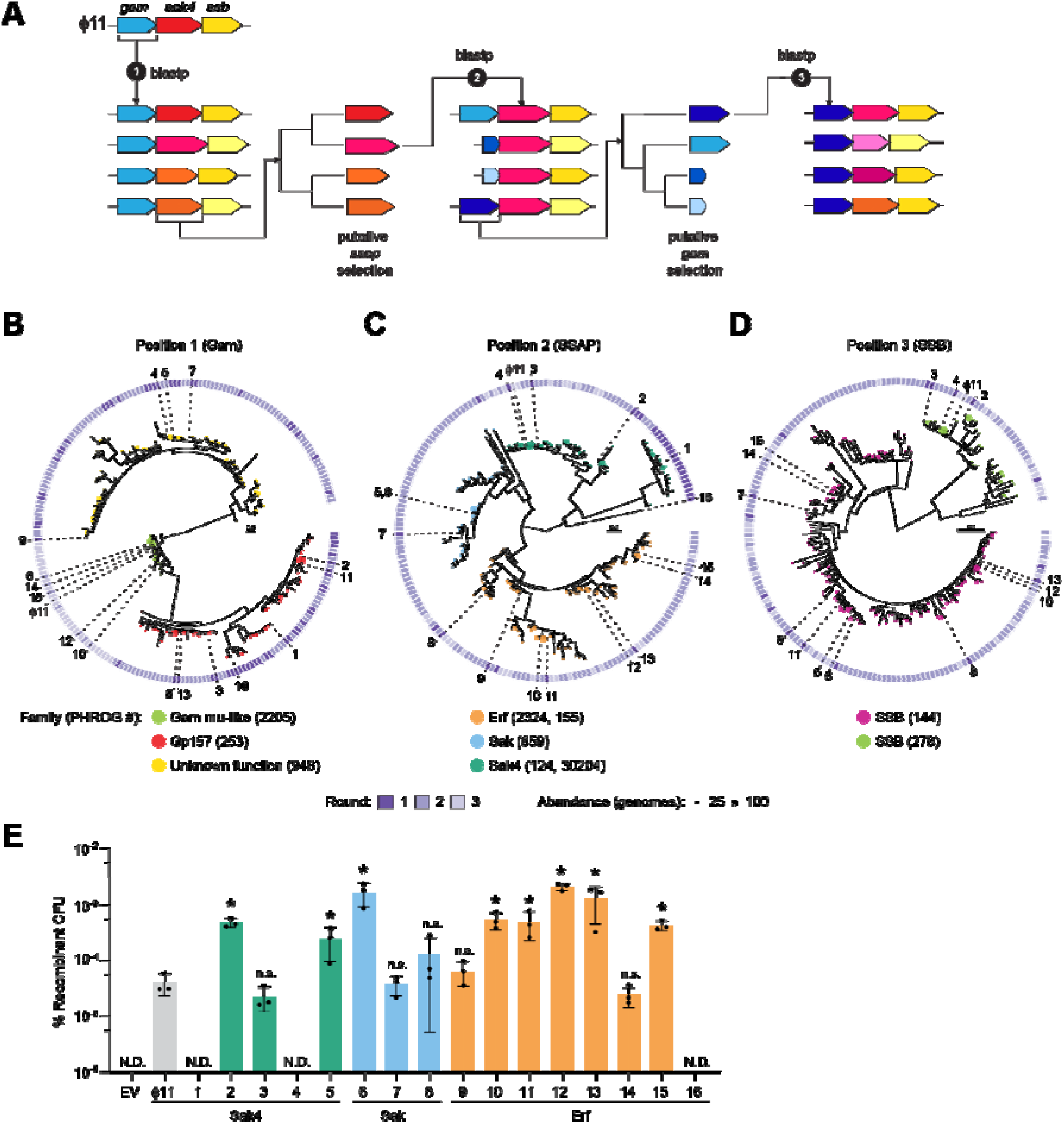
Mining of *Staphylococcus* genomes identifies more efficient phage recombination cassettes. **(A)** Schematic of the iterative, synteny-guided search strategy. (1)□11 Gam from the SapSak4 system was queried in a protein–protein BLAST. Putative SSAP/recombinases and SSBs were extracted downstream from a representative genome for each BLAST hit, and seven putative SSAP/recombinases were selected for round 2. (2) The seven recombinases were used to perform the same search as in (1) and the putative Gams and SSBs neighboring each BLAST hit were extracted and six putative Gam sequences were selected for round 3. (3) Same as in (1) using the six sequences from round 2. Ten putative SSAP/recombinases were selected at the end of round 3. Sixteen 3-gene loci harboring the putative Gam and SSAP/recombinases (Supplementary Figure S2) selected in the iterative search were cloned under IPTG-inducible promoters in an *S. aureus* vector. **(B)** Maximum-likelihood phylogenetic tree of the putative Gam proteins identified during the iterative blastp search in **(A)**. Sequences clustered at 90% identity are shown together with the sixteen candidates selected and 11 Gam. The end of each branch is colored by PHROG family and the tip size represents the number of genomes within each 90%-identity group. Outer track is colored by which blastp round a genome from that 90%-identity cluster was first observed. **(C)** Same as in (B), but for the putative SSAP/recombinases. (D) Same as in **(B)**, but for the putative SSBs (candidates 1 and 16 are absent because the third gene in their locus is not a putative SSB). **(E)** Percent of recombinant CFU following electroporation of a dsDNA template containing an antibiotic-resistance gene into *S. aureus* RN4220 harboring a vector with one of the sixteen candidates, the SapSak4 recombination cassette (labeled□11), or an empty vector. Means of three biological replicates ± SD are reported. *, adjusted P < 0.05 (all marked comparisons ≤ 0.0078); n.s., not significant (adjusted P > 0.05); ordinary one-way ANOVA of log10-transformed data with Dunnett’s multiple-comparisons test against the SapSak4 cassette. N.D., no recombinants detected; the empty vector and candidates 1, 4, and 16 were excluded from the statistical analysis.

Many of the recovered recombinases were too divergent to classify by sequence alone. We therefore annotated each protein using a combination of predicted structure and phage-specific profile searches (53–55), comparing predicted folds against reference SSAPs/recombinases (56). Structure-based classification identified many recombinases that sequence searches had missed and resolved the collection into three SSAP/recombinase families: the Sak4 family (belonging to the RAD51-superfamily), which contains the□11 recombinase, SapSak4, and the Erf and Sak families (RAD52-superfamily) (40,41). Notably, the□11-type Sak4 family was the least abundant of the three, present in only 9% of cassettes, whereas Erf was by far the most common at 48%. To map this diversity and locate the SapSak4 system from □11 within it, we constructed phylogenies for each position of the cassette, Gam, recombinase, and SSB (Figure 2B–D). SapSak4 fell within a single, well-populated Sak4 clade, while the Erf and Sak families formed separate and equally diverse groups, indicating that staphylococcal phages deploy several unrelated recombinases/SSAPs within the same conserved position (Figure 2C).

The phylogenies of the two flanking genes revealed contrasting relationships. At the Gam position, the phylogeny separated a clade of Mu Gam–like proteins, which included the □11 Gam-like proteins, a set of proteins that aligns to the siphovirus Gp157 protein family, and a distinct, much smaller (∼74 aa) DUF2483-family protein of unknown function (Figure 2B). A comparable set of genes upstream of recombinases has previously been reported in *Firmicutes* phages (52). Neither group was specific to a recombinase: the same families recurred across unrelated recombinase superfamilies, consistent with the Gam acting as a generic, interchangeable protectant of free DNA ends rather than a cognate partner of the recombinase. In contrast, the SSB position revealed a distinct pattern. Here the proteins resolved into two well-separated families (Figure 2D) that segregated cleanly by recombinase superfamily: across all recovered cassettes the RAD52-superfamily Erf and Sak recombinases were paired almost exclusively with one SSB family (PHROG 44), whereas the RAD51-superfamily Sak4 recombinases, including that of □11, never carried this family and instead used a different SSB subset (PHROG 378). This mutually exclusive pairing indicates that each recombinase and its SSB are cognate partners that have coevolved within phage recombination cassettes.

Guided by these phylogenies, we selected a representative panel of 16 candidate cassettes spanning the three SSAP families, a potential unassigned recombinase, and the range of Gam variants (Supplementary Figure S2), and synthesized them for functional testing in *S. aureus.* To test whether the predicted cassettes could mediate recombineering, we cloned each system into the inducible *S. aureus* vector and assayed dsDNA editing at the *sau1hsdR* locus of *S. aureus* RN4220 (Figure 2E). Thirteen of the sixteen potential candidates were able to mediate recombineering, confirming that our synteny-guided search was able to reliably recover functional recombination systems. Notably, ten of these systems performed better than the original SapSak4 recombination system from□11. Activity varied widely, both between and within families. The Erf family was the most active and the most reproducible: the strongest Erf systems edited at rates up to three logs higher than the SapSak4 system. Sak4 systems, the family to which □11’s SapSak4 itself belongs, were more variable; only one (candidate 2) approached the best Erf cassettes, whereas two others produced either low or no detectable editing.

The three cassettes that failed to edit each lacked a functional version of a different component, reinforcing the requirement for all three. Two were missing a recognizable SSB: in candidate 1 the SSB position was occupied by an HNH endonuclease, and candidate 16, which also carried a protein that could not be assigned to any SSAP family, harbored a small 50 aa protein of unknown function. The third, candidate 4, carried an intact cognate SSB and a Sak4 recombinase of the same family as SapSak4, but combined them with a DUF2483-family protein of unknown function in the *gam* position. Among closely related Sak4 cassettes differing essentially only in their *gam* position, editing was weak but detectable with a Gam (□11, candidate 3) and fell below detection with the small Gam (candidate 4). The same trend held in a matched comparison of identical Sak recombinases, where substituting the DUF2483 protein for the Gam or Gp157 reduced editing roughly sevenfold (candidates 5 and 6). These observations fit our earlier findings for the SapSak4 system, where the Gam is not strictly required but greatly increases the efficiency of editing.

In summary, our results showed that the most abundant family in our survey, Erf, mediated the most efficient recombineering. However, the recombinase/SSAP family identity predicts but does not fully determine activity, as efficiencies differed among members of the same family, implicating cassette-specific features beyond the recombinase/SSAP group. We therefore moved forward with the most active and independently reproducible Erf candidate (candidate 13), designated SppErf (*S. pseudintermedius* prophage Erf) as the basis for a recombineering tool.

### The SppErf cassette enables efficient dsDNA and ssDNA recombineering in *S. aureus*

The SppErf system derives from a prophage carried by *S. pseudintermedius* strain MAD568, whose closest cultured relative is *Staphylococcus* phage SP120 (genus *Coventryvirus*) (Figure 3A and Supplementary Figure S2). The locus pairs a RAD52-superfamily Erf annealase with Gp157 upstream and a downstream cognate SSB. Gp157 is a hypothetical protein whose homologs have been assigned to the replication module of *Streptococcus thermophilus* phages (57). Although Gp157 shares no detectable amino acid similarity with the Gam protein of *E. coli* phage Mu, structural prediction revealed a closely matching fold to the solved structure of the Mu-like Gam in *Desulfovibrio vulgaris* (Figure 3B). This suggests that Gp157 also protects free dsDNA ends, analogous to 11’s Gam-like protein from the SapSak4 system. This configuration is not confined to prophages: in an unbiased survey independent of our synteny-based search, ∼11% of the 1,018 staphylococcal phage genomes in INPHARED encode a complete Gam–SSAP–SSB cassette with an Erf annealase.

**Figure 3.**
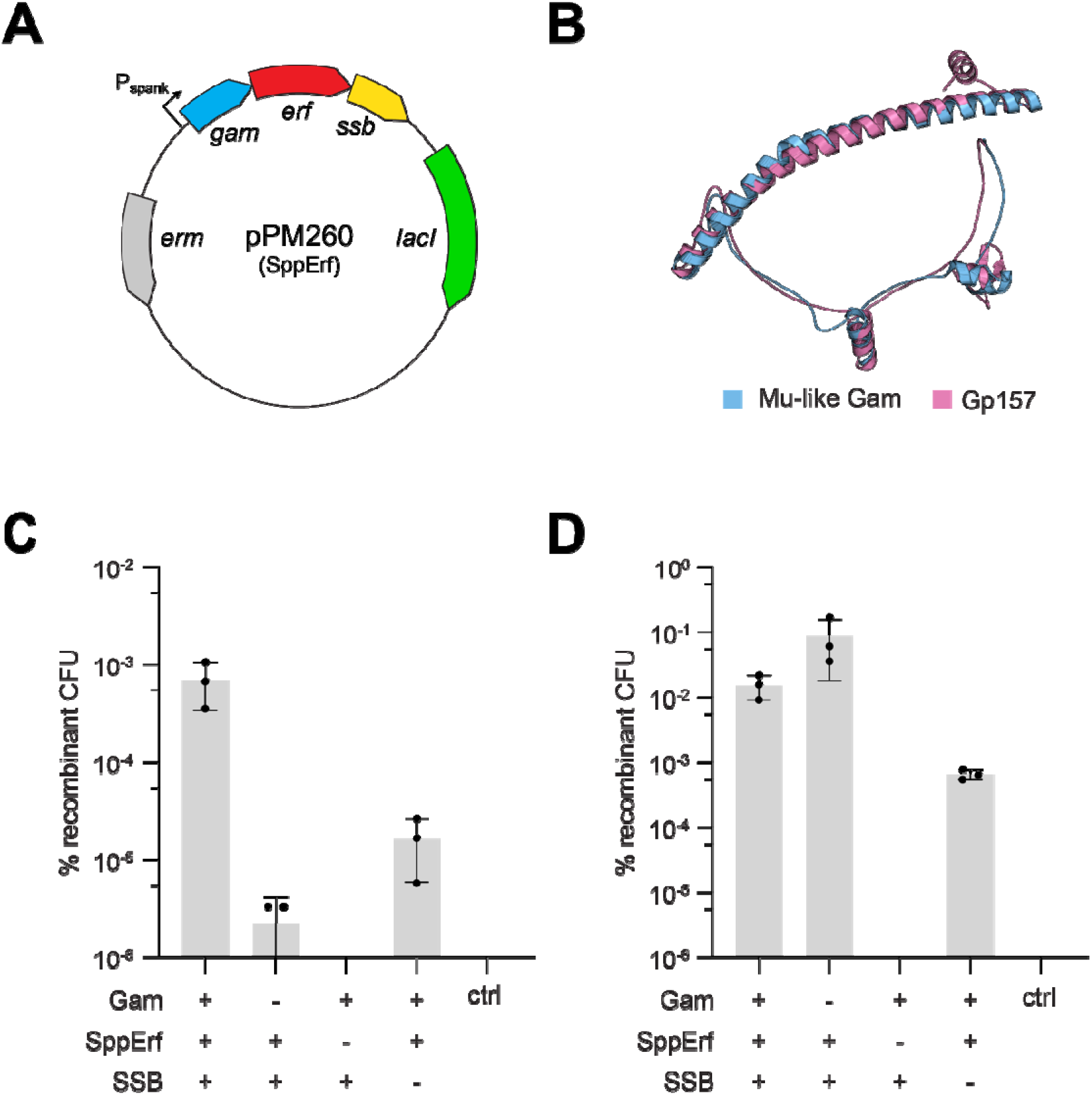
The SppErf cassette (candidate 13) mediates efficient dsDNA and ssDNA recombineering in *S. aureus*. **(A)** Schematic of the recombineering vector pPM260. The plasmid contains an IPTG-inducible operon with *S. pseudintermedius* MAD568 *gam*, *erf*, and *ssb* together with an erm selection marker and lacI. **(B)** Superimposition of the Mu-like Gam structure 2P2U (blue) and the predicted structure of Gp157 (orange). **(C)** Percent of recombinant CFU following electroporation of a dsDNA template containing a chloramphenicol-resistance cassette flanked with homology to the *sau1hsdR* locus into *S. aureus* RN4220 harboring pPM260 or derivatives lacking *gam*, *erf*, *ssb*, or all three genes (ctrl, an empty vector). Means of three biological replicates ± SD are reported. **(D)** Same as in **(C)**, but following electroporation of *S. aureus* sPM05 with a single-stranded oligonucleotide template for repair of a chromosomal chloramphenicol-resistance cassette or water (ctrl). Means of three biological replicates ± SD are reported.

We first established the conditions under which the SppErf system operates most efficiently. Extending induction to 45 min and increasing the recombination template to 2 µg each improved editing (Supplementary Figure S3A and B). Additionally, edited cells were recovered with homology arms as short as 60 bp, although maximum editing was observed at 2 kb (Supplementary Figure S3C). With this optimized protocol we tested the genetic requirements for dsDNA recombineering with the SppErf system. Editing was highest when all three proteins were expressed (Figure 3C). As in the□11 cassette, removal of the Gam reduced efficiency sharply; in contrast to□11, editing remained detectable in the absence of the SSB, likely reflecting the higher overall efficiency of this system (Figure 3C). Together, these results define an optimized protocol and confirm that all three components are required for maximal efficiency, but unlike Φ11, the SppErf cassette retains detectable dsDNA editing without the SSB

We next asked whether the SppErf system could also recombine ssDNA. We constructed a reporter strain carrying a chloramphenicol-resistance gene inactivated by non-functional start codon and attempted to restore resistance through recombineering of a ssDNA oligonucleotide. (Supplementary Figure S4A). As for dsDNA, we first optimized the conditions for ssDNA recombineering in *S. aureus*. In common with other recombineering systems, oligonucleotides targeting the lagging strand edited at higher rates than those targeting the leading strand (Supplementary Figure S4B). Incorporating additional silent mismatches into the oligonucleotide has been reported to evade the host mismatch repair machinery and raise recombination rates (58,59), but we observed no positive effect (Supplementary Figure S4B). Oligonucleotide length over the range of 79–99 nucleotides likewise had little effect on editing efficiency (Supplementary Figure S4C), whereas increasing oligonucleotide concentration, or adding a 5′ phosphorothioate linkage (60), each improved efficiency (Supplementary Figure S4D and E). With this optimized protocol we edited just under 1% of all cells, a ceiling likely set by the transformation efficiency of *S. aureus* (Figure 3D). In contrast to dsDNA recombineering, SppErf was the only component strictly required: removal of the Gam had no effect, as expected for a template that presents no free duplex ends, while removal of the SSB reduced editing efficiency by approximately two orders of magnitude (Figure 3D).

### Coupling SppErf recombineering with Cre–*lox* excision for markerless, serial editing

Because selection for dsDNA recombineering relies on inserting an antibiotic-resistance cassette at the target locus, the edited strain retains a resistance marker and a large genomic insertion, which complicates serial editing. To remove the marker after selection, we built a derivative of the SppErf recombineering vector carrying an *S. aureus* codon-optimized *cre* recombinase under an anhydrotetracycline (aTc)-inducible promoter (Figure 4A). In this two-step scheme, a donor in which the chloramphenicol-resistance (*cat*) cassette is flanked by *loxP* sites is first recombineered into the target locus under IPTG induction and selected; subsequent induction of Cre excises the cassette, leaving a single *loxP* site in place of the marker (Figure 4B).

**Figure 4.**
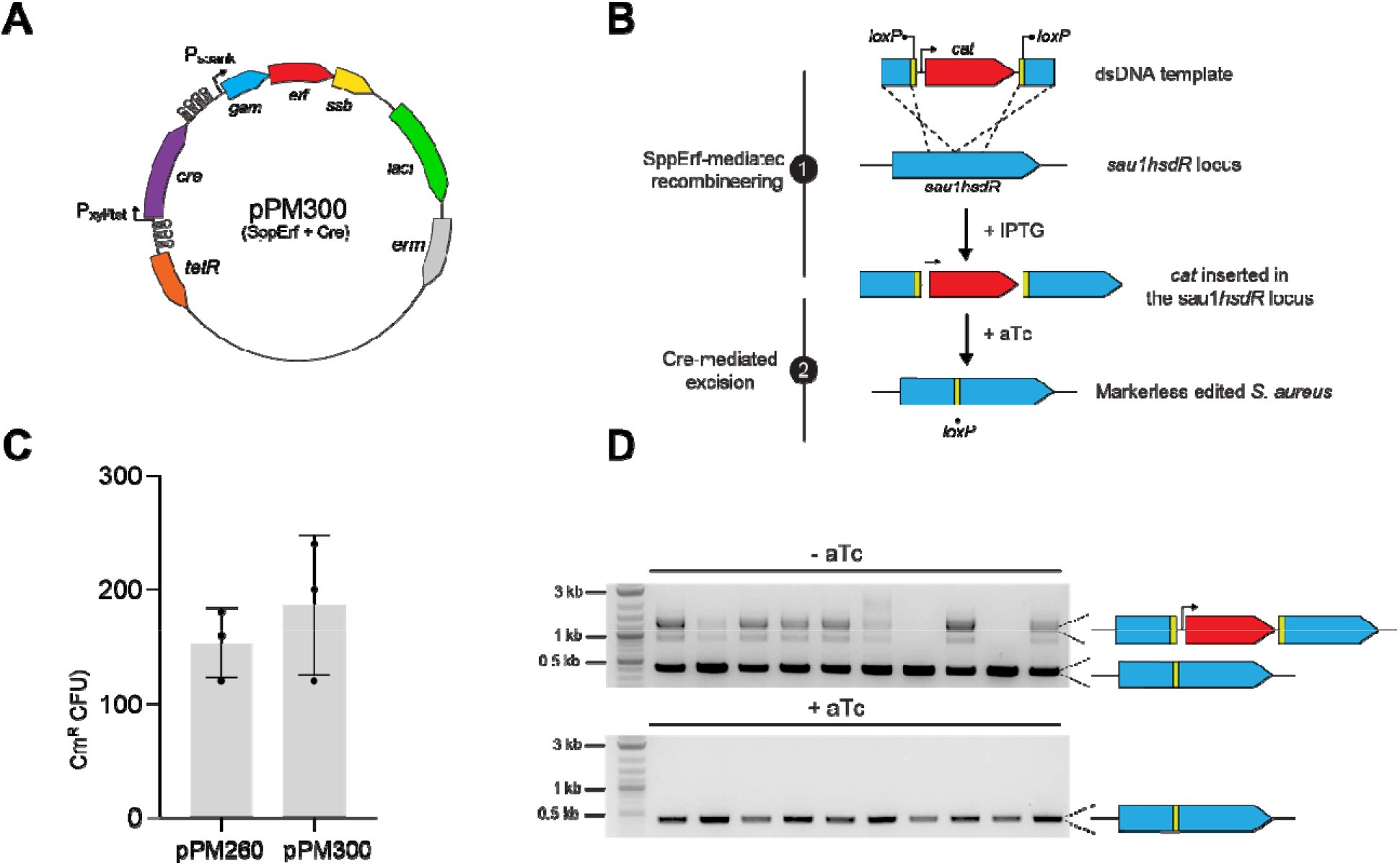
Coupling SppErf recombineering with Cre-mediated excision enables markerless editing in *S. aureus*. **(A)** Schematic of the vector pPM300, a derivative of pPM260, harboring the IPTG-inducible 3-gene recombination cassette from *S. pseudintermedius* MAD568 together with an *S. aureus* codon-optimized cre recombinase under an anhydrotetracycline (aTc)-inducible promoter. **(B)** Schematic of a 2-step markerless editing strategy. (1) IPTG induction of the recombineering cassette together with electroporation of a dsDNA template harboring a floxed chloramphenicol-resistance cassette flanked with homology to the *sau1hsdR* locus, resulting in insertion of the cassette in the genome. (2) Cre is aTc-induced in recombinant colonies to mediate excision of the chloramphenicol-resistance cassette. **(C)** Chloramphenicol-resistant colonies following electroporation of *S. aureus* RN4220 with a dsDNA template encoding a floxed chloramphenicol-resistance cassette flanked with homology to the *sau1hsdR* locus. Electroporated *S. aureus* harbored either the recombineering cassette from S. pseudintermedius MAD568 alone (pPM260) or the cassette together with aTc-inducible cre (pPM300). Means of three biological replicates ± SD are reported. **(D)** Agarose gel electrophoresis of PCR products obtained after amplification of the *sau1hsdR* locus from 10 single colonies of a pPM300-harboring recombinant culture from (C), treated with (bottom) or without (top) 125 ng/mL aTc for five hours.

One concern with this design is that basal *cre* expression from the vector might interfere with the initial recombination step, either by reducing recombination or by excising the marker before selection. To test this, we compared the dsDNA editing efficiency of the Cre-encoding vector with that of the original SppErf recombineering vector. Neither the presence of *cre* on the vector nor the *loxP* sites on the donor substantially reduced recombination efficiency (Figure 4C). We then tested excision of the integrated cassette. An edited clone carrying the new vector was grown with or without aTc, and the edited locus was examined by PCR. Partial excision was detected in the absence of aTc, consistent with some basal Cre activity, whereas the ∼1 kb *cat* cassette was completely lost in all induced colonies (Figure 4D and Supplementary Figure S5A). Combined with IPTG-induced curing of the recombineering plasmid (Supplementary Figure S5B), this yields a markerless, plasmid-free edited strain and enables iterative rounds of editing in *S. aureus*. In practice, the entire procedure, from electroporation of the donor to a verified, markerless edit, is completed in four days: recovery and selection of the initial edit (days 1–2), aTc-induced Cre excision of the resistance cassette (day 3), and PCR confirmation of the markerless locus together with IPTG-mediated curing of the recombineering plasmid (day 4).

### The SppErf recombineering system functions across bacterial species

The Erf cassette we identified addresses two of the major obstacles to portability encountered with isolated SSAPs: (i) it encodes its own Gam, which protects linear DNA by binding free duplex ends rather than by contacting a specific host nuclease, and (ii) it encodes its own SSB. Therefore, the SppErf system should function independently of the host and could be used for recombineering in diverse bacterial species. To test this, we first introduced the SppErf system into a panel of clinical *S. aureus* isolates, which are typically refractory to editing because they retain intact restriction–modification systems and transform poorly. Editing was achieved in every strain tested (Figure 5A). We then constructed inducible expression vectors for *Streptococcus pyogenes*, *Enterococcus faecium*, *Francisella novicida*, and *Escherichia coli*. This system supported editing in all four organisms, albeit to varying degrees, and was most efficient in *S. pyogenes* (Figure 5B). Two factors appear to govern efficiency. First, editing declined with increasing evolutionary distance from *S. aureus*: rates in *E. coli* and *F. novicida* were substantially lower than in *S. aureus*. Second, the transformation efficiency of the recombineering construct presents a major bottleneck; in *E. faecium*, where transformation is difficult, editing rates were the lowest despite its close relationship to *S. aureus*. Despite these caveats, these results demonstrate the portability of the Erf recombineering system across phylogenetically distant bacteria.

**Figure 5.**
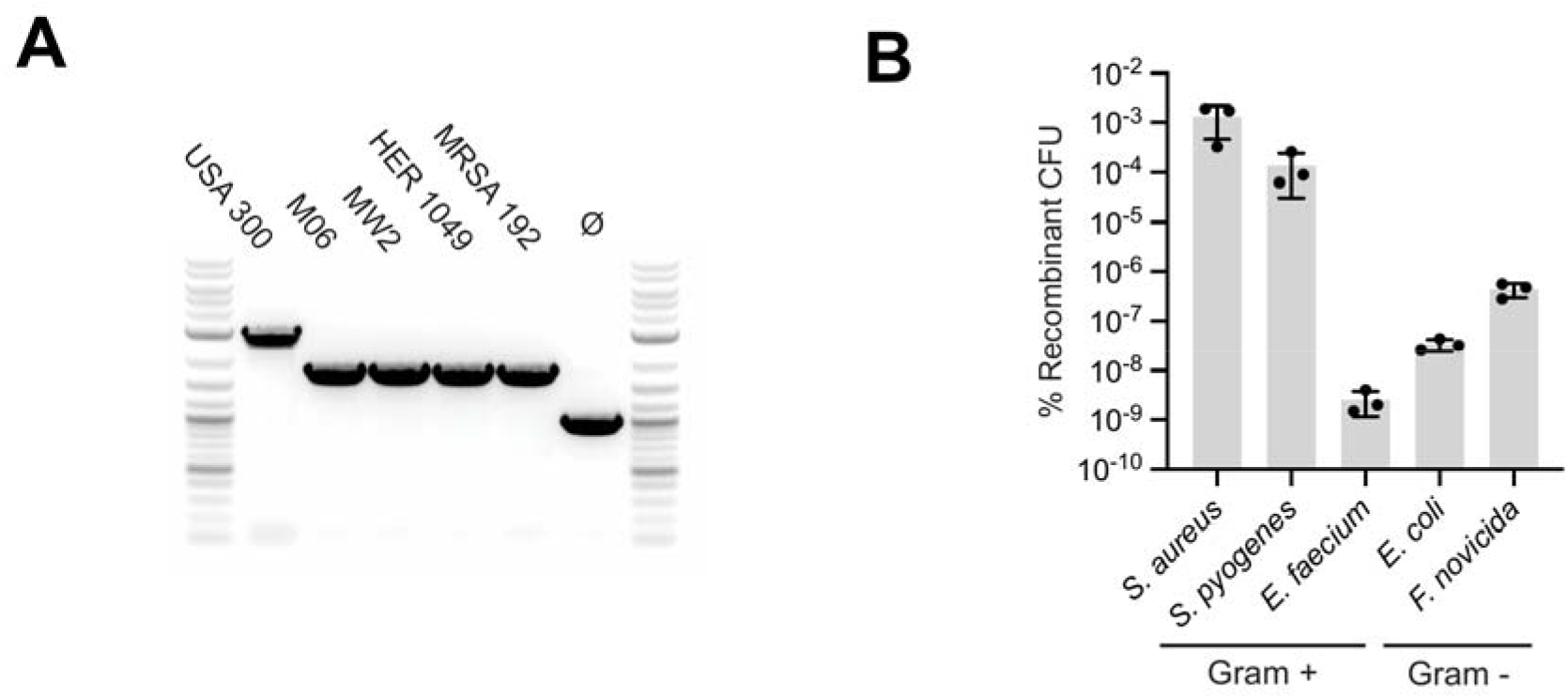
The SppErf recombineering system edits clinical *S. aureus* isolates and diverse bacterial species. **(A)** PCR of the edited *sau1hsdR* locus in clinical *S. aureus* isolates showing spectinomycin (USA300) or chloramphenicol insertion (M06, MW2, HER1049, and MRSA192) with the Erf recombineering vector following electroporation of a dsDNA substrate. Ø, unedited control. A representative result is shown. **(B)** Percent of edited cells following recombineering with species-specific inducible recombineering expression vectors in the Gram-positive *S. aureus*, *S. pyogenes*, and *E. faecium* and the Gram-negative *E. coli* and *F. novicida*. Means of three biological replicates ± SD are reported.

## Discussion

Efforts to extend recombineering to new bacteria have largely focused on the recombinase, treating the annealase as the unit of transfer and leaving nuclease protection and ssDNA loading to the host. Here we describe a class of phage recombination cassettes in which all three activities are encoded together and show that this complete unit mediates recombineering across phylogenetically distant bacteria. The SppErf cassette we characterized encodes a Mu Gam-like protein that protects linear dsDNA from host nucleases, an Erf-family SSAP, and a cognate SSB. In *S. aureus* the SppErf system supports markerless editing on a four-day timescale and ssDNA editing in close to 1% of all cells. Because the SppErf cassette supplies both nuclease protection and ssDNA loading itself, it does not depend on host-specific interactions, and we found it to be functional in *S. pyogenes*, *E. faecium*, *F. novicida*, and *E. coli*.

We show that the SSB encoded by the cassette is essential for function, and that SSB subgroups associate tightly with their cognate SSAP/recombinase. This is consistent with previous work showing that a properly paired SSB is required for phage function in *S. aureus* (42), and with the demonstration that lambda Red depends on the host *E. coli* SSB (11). Together these observations suggest a broader pattern in which SSAP/recombinases engage specific SSBs, and in which that pairing is required for activity. How the interaction is achieved at the molecular level remains to be determined.

Recent work has substantially broadened the reach of recombineering by expanding the repertoire of single-strand annealing proteins available for a given host. High-throughput screens identified individual annealases with markedly improved activity in *E. coli* and *Pseudomonas* (32) and a pooled library of 227 SSAPs was recently shown to support efficient editing, including double-stranded and multiplexed editing in *S. aureus*, across bacteria from three divergent classes (33). These approaches share a common strategy: they search a collection of isolated annealases for a variant that functions in the target organism, while still relying on that organism’s own single-stranded DNA-binding protein and nuclease-protection machinery (11). Our results describe a complementary strategy. Rather than screening many annealases per host, we transfer a single cassette that supplies the accessory activities itself, so it behaves as a defined, portable unit requiring no host-specific optimization. We therefore view self-sufficient operons and diverse annealase libraries as complementary approaches toward the goal of universal bacterial genome editing.

While functional across a wide range of species, the SppErf cassette loses efficiency in distantly related hosts such as *E. coli*, even though these are among the most readily transformable organisms tested. Several explanations are possible. Translation of the cassette may be less efficient outside its native host. Alternatively, the SppErf system may depend on host factors we have not yet identified, of which the host exonuclease is a prime candidate. For a dsDNA substrate, one strand must be resected before annealing can occur; in lambda Red this is carried out by Exo, and in RecET by RecE. Our cassette encodes no exonuclease, and so presumably relies on host machinery, RecBCD, AddAB, RecJ, or their equivalents. Importantly, the reliance on host resection activity does not require specific interactions with the recombineering machinery and presumably many non-orthologous enzymes across the bacterial kingdom should be able to generate the 3′ overhangs required for recombination. A generic enzymatic requirement may be more readily satisfied in a new host than a specific interaction surface, which may explain why the cassette retains activity in distant hosts while losing efficiency.

The Gam-like protein may contribute to this tolerance in a similar manner. Mu Gam binds and occludes free dsDNA ends, rather than binding to and inhibiting host nucleases, as is the case for lambda Gam with RecBCD in *E. coli*. A protein that acts on the substrate rather than on a host enzyme has no interaction to maintain as it moves between species, and should therefore protect incoming dsDNA recombination templates regardless of the nature of host exonucleases.

Finally, these results suggest that the limits on recombineering portability are set not by the strand-annealing reaction itself, but by the requirement that the annealase engage *host* nucleases and ssDNA-binding proteins. Encoding nuclease protection and ssDNA loading in cis removes that requirement: the cassette carries its own interaction partners and no longer depends on compatible host factors. This opens the possibility of designing cassettes for organisms in which no recombineering system has been established. Further work is needed to determine whether resection can also be encoded within the cassette, and how far the approach extends, including, speculatively, to eukaryotic systems where Rad52-family annealases perform analogous reactions. Overall, these results reframe the unit of transfer in recombineering and establish a framework for building recombination systems in hosts that remain genetically intractable.

## Acknowledgments

We thank Jakob Træland Rostøl and Howard Hang for sharing reagent and for helpful discussions.

## Author contributions

Conceptualization: PM, LM, AV

Investigation: PM, MA, MB, VC, JZ, KS, LM, AV

Funding acquisition and supervision: LM, AV

Writing – original draft: PM, AV

Writing – review & editing: PM, MA, MB, VC, JZ, KS, LM, AV

## Data and materials availability

All data, code, and materials used in this study are available upon request. All custom code has been deposited on GitHub (https://github.com/andrew-varble/Varble-Lab).

## Supplementary Materials

### Supplementary Figures

**Figure S1.**
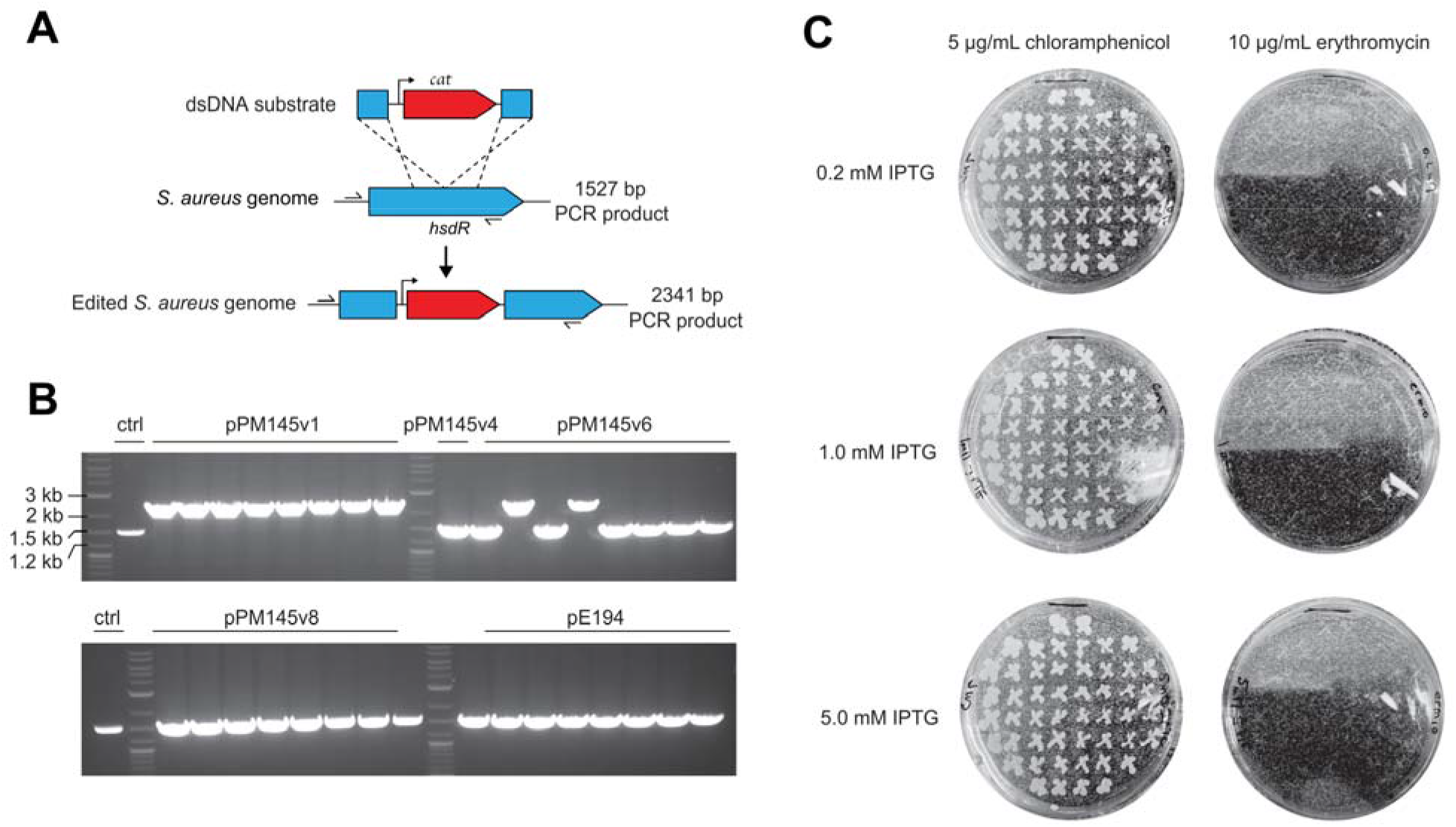
SapSak4 recombineering: confirmation of editing and plasmid curing. **(A)** Schematic of the dsDNA substrate and the *sau1hsdR* locus, indicating the expected colony-PCR products before (1,527 bp) and after (2,341 bp) editing. **(B)** Colony PCR confirming editing in *S. aureus* RN4220 carrying pPM145 variants (pPM145v1, v4, v6, v8) or controls (ctrl, pE194), resolved on an agarose gel. **(C)** Plasmid curing. *S. aureus* carrying the recombineering plasmid was grown with 0.2, 1.0, or 5.0 mM IPTG and patch-plated on chloramphenicol (left) or erythromycin (right); loss of erythromycin resistance indicates loss of the recombineering plasmid.

**Figure S2.**
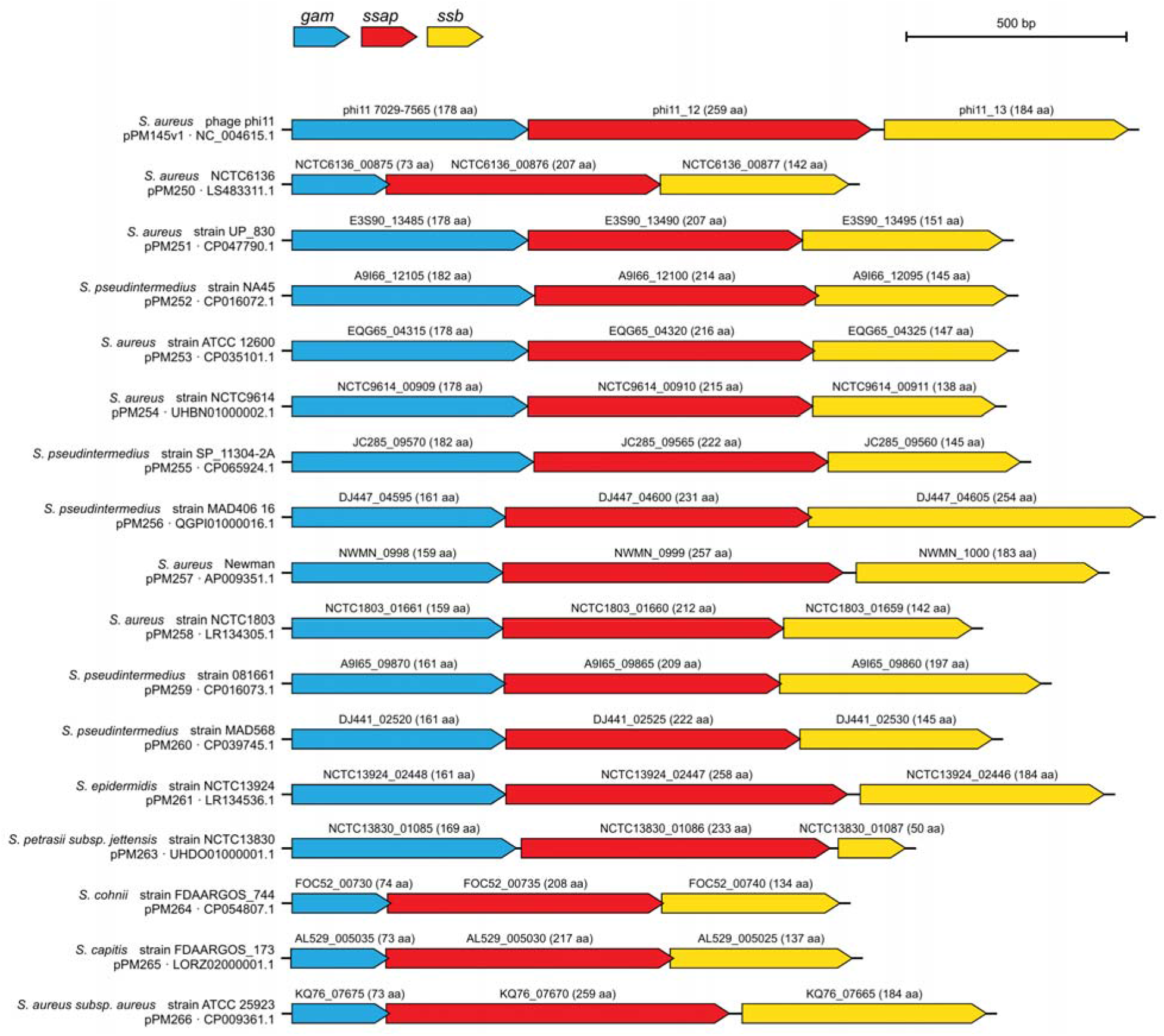
Gene maps of the candidate recombination cassettes selected for testing. Gene-neighborhood maps, drawn to scale, of the SapSak4 reference cassette from□11 (pPM145v1) and the 16 candidate cassettes selected from the synteny-guided search (Figure 2). For each cassette the *gam* (blue), *SSAP/recombinase* (red), and *ssb* (yellow) genes are shown, annotated with the source organism, plasmid, protein accession, and length (aa). Scale bar, 500 bp.

**Figure S3.**
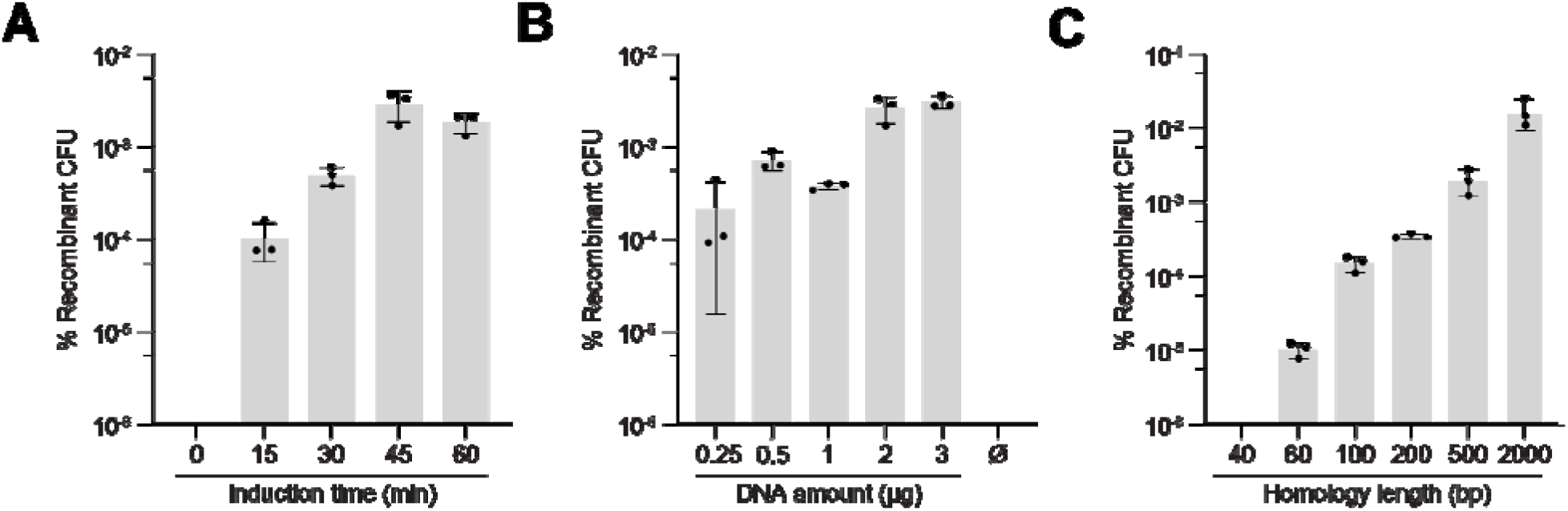
Optimization of dsDNA recombineering with the SppErf system. Percent of edited cells in *S. aureus* RN4220 harboring pPM260 following electroporation of a dsDNA substrate, varying: **(A)** IPTG induction time before electroporation (0–60 min); **(B)** amount of dsDNA substrate (0.25–3 µg; Ø, no DNA); **(C)** length of the homology arms flanking the resistance cassette (40–2,000 bp). Means of three biological replicates ± SD are reported.

**Figure S4.**
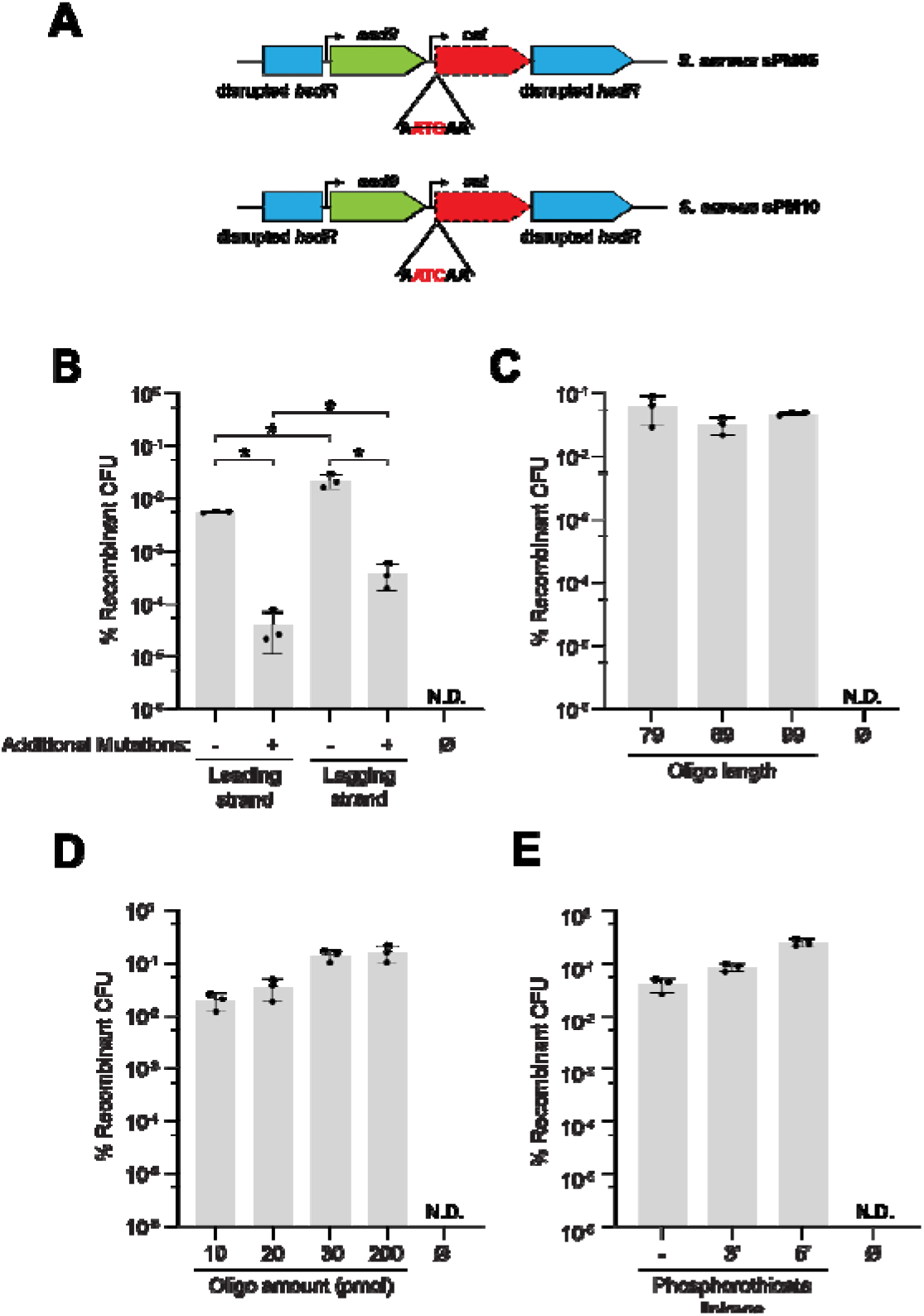
Optimization of ssDNA recombineering with the SppErf system. **(A)** Schematic of the ssDNA reporter strains *S. aureus* sPM05 and sPM10, in which a chloramphenicol-resistance gene (*cat*) is inactivated by a point mutation (sPM10) or the removal of the start codon (sPM05). This cassette is integrated at the *sau1hsdR* locus alongside a spectinomycin-resistance gene (*aad9*); recombineering with an oligonucleotide restores *cat*. In all panels, means of three replicates ± SD are reported. **(B)** Percent of recombinant CFU following electroporation of *S. aureus* sPM05 with oligonucleotides targeting the leading or lagging strand, each with or without additional silent mismatches intended to evade mismatch repair; Ø, no oligo; *, adjusted P < 0.05; ordinary two-way ANOVA of log10-transformed data with Tukey’s multiple-comparisons test (adjusted P: leading strand, − vs + mismatches, <0.0001; lagging strand, − vs + mismatches, <0.0001; − mismatches, leading vs lagging, 0.0297; + mismatches, leading vs lagging, 0.0015). N.D., no recombinants detected; the no-oligonucleotide control (Ø) was excluded from the statistical analysis. **(C)** As in **(B),** but in *S. aureus* sPM10, using lagging-strand oligonucleotides without additional mismatches of varying length (79, 89, 99 nt). **(D)** As in **(C)**, but using *S. aureus* sPM05, varying oligonucleotide amount (10– 200 pmol). **(E)** As in **(C)**, comparing unmodified oligonucleotides with those carrying phosphorothioate linkages at the 3′ or 5′ terminus.

**Figure S5.**
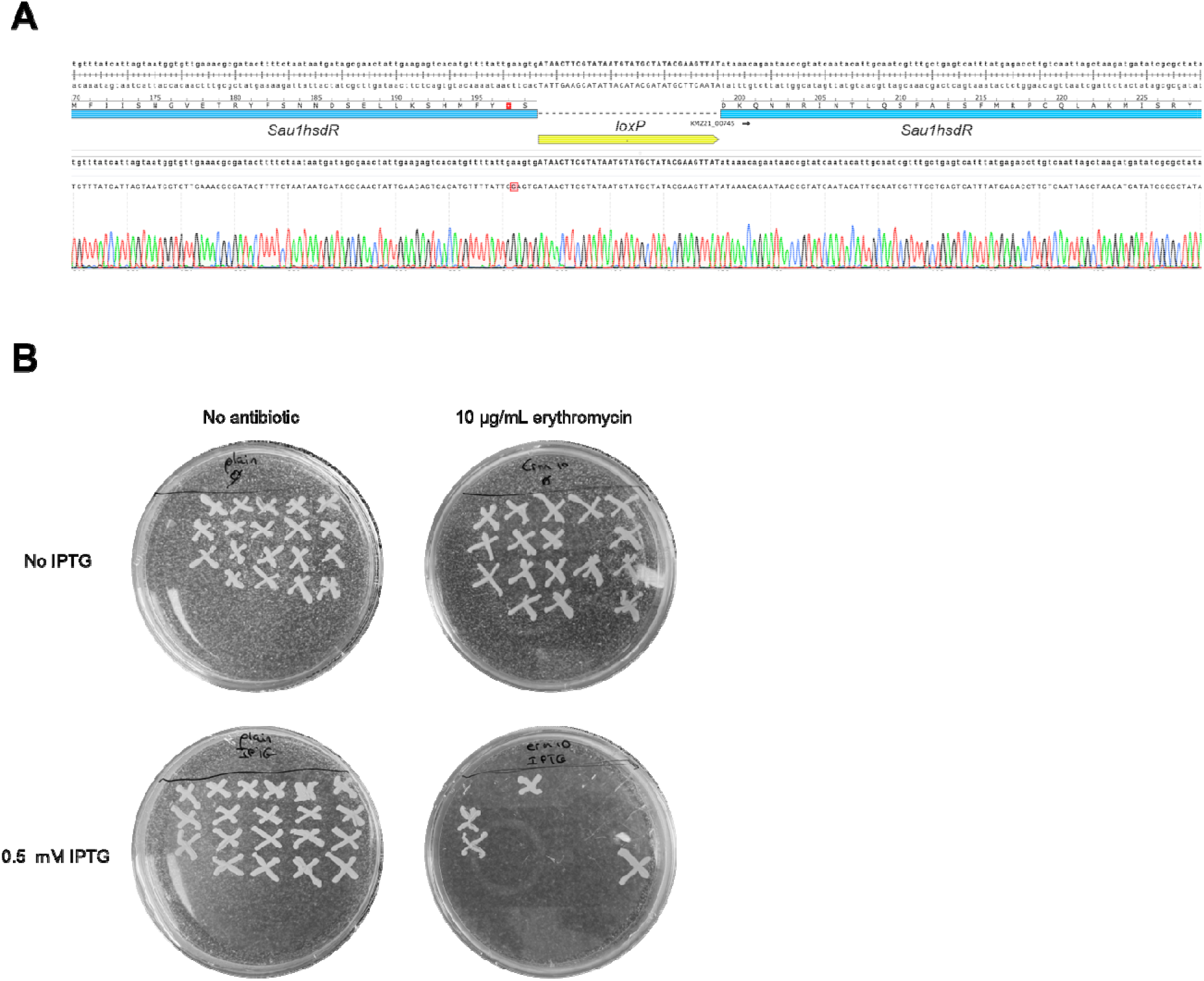
Cre-mediated excision of the resistance cassette and curing of the SppErf recombineering plasmid. **(A)** Chromatogram of PCR products obtained after amplification of the *sau1hsdR* locus from single colonies of pPM300-harboring recombinants, confirming complete loss of the ∼1 kb floxed chloramphenicol-resistance cassette in induced colonies. **(B)** Plasmid curing. *S. aureus* carrying the recombineering plasmid pPM300 was grown with or without 0.5 mM IPTG and patch-plated on no antibiotic (left) or erythromycin (right); loss of erythromycin resistance indicates loss of the recombineering plasmid.

### Materials and Methods

#### Bacterial strains and growth conditions

Culturing of *S. aureus* was carried out in brain–heart infusion (BHI) medium at 37 °C. Liquid cultures (5 mL) were grown in 15 mL conical tubes at 37 °C with shaking at 220 rpm. For antibiotic selection, *S. aureus* was grown in the presence of chloramphenicol at 10 µg/mL (pC194-based vector selection (61)) or 5 µg/mL (chromosomal selection), spectinomycin at 250 µg/mL (plasmid selection) or 125 µg/mL (chromosomal selection), and erythromycin at 10 µg/mL for pE194-based vectors (62).

Culturing of *S. pyogenes* was carried out in BHI medium at 37 °C. Liquid cultures (10 mL) were grown in 15 mL conical tubes at 37 °C without agitation. For antibiotic selection, *S. pyogenes* was grown in the presence of 5 µg/mL chloramphenicol and 50 µg/mL spectinomycin.

Culturing of *E. coli* was carried out in Luria broth (LB) at 37 °C. Liquid cultures (5 mL) were grown in 15 mL conical tubes at 37 °C with shaking at 220 rpm. For antibiotic selection, *E. coli* was grown in the presence of 30 µg/mL chloramphenicol. For arabinose, arabinose was added to a final concentration of 2 mM.

Culturing of *E. faecium* was carried out in BHI medium at 37 °C. For antibiotic selection*, E. faecium* was grown in the presence of 10 µg/mL chloramphenicol, 50 µg/mL erythromycin, or 250 µg/mL spectinomycin. For thymidine auxotrophy assays, *E. faecium* was grown on MM9YEG agar (1× M9 salts, 0.25% yeast extract, 0.5% glucose, 1.5% agar) supplemented with 40 µg/mL thymidine and chloramphenicol when appropriate.

Culturing of *F. novicida* was carried out in BHI medium at 37 °C. Liquid cultures were grown aerobically at 37 °C in BHI broth supplemented with 0.1% L-cysteine. For antibiotic selection, *F. novicida* was grown in the presence of 50 µg/mL ampicillin, 15 µg/mL kanamycin for strains carrying pFNMB2-based plasmids (63), or 30 µg/mL erythromycin for recombineered strains encoding the ermC methylase. For aTc induction of recombineering cassettes, aTc was added to a final concentration of 100 ng/mL.

#### Bacterial transformation for recombineering

Electrocompetent *S. aureus* cells were prepared by diluting overnight cultures 1:100 in BHI + 0.5 M sorbitol (BHIS) and cultured at 37 °C with shaking at 220 rpm. At OD_600_ = 0.3–0.4, isopropyl β-D-thiogalactopyranoside (IPTG) was added to a final concentration of 1 mM and the cultures were incubated further until OD_600_ = 0.8. Following incubation, cells were pelleted at 4,300 rpm, washed twice by resuspension in the original culture volume of 0.5 M sucrose, then pelleted again and resuspended in 0.5 M sucrose to OD_600_ = 25. For each electroporation, 100 µL of the resuspended culture was mixed with DNA in a 2 mm cuvette and electroporated using a Gene Pulser Xcell instrument (Bio-Rad) with the following settings: 2,900 V, 25 µF, 100 Ω, 2 mm.

Electrocompetent *S. pyogenes* cells harboring pAV488 were prepared by diluting overnight cultures of the pAV488-containing strain 1:50 in 50 mL BHI containing 5 µg/mL chloramphenicol and cultured at 37 °C without agitation. At OD_600_ = 0.15, isopropyl β-D-thiogalactopyranoside (IPTG) was added to a final concentration of 1 mM and the cultures were incubated further until OD_600_ = 0.3. Following incubation, cells were pelleted at 4,300 rpm for 15 min at 4 °C, washed twice in 50 mL cold 10% glycerol (pelleting at 12,000 rpm for 15 min at 4 °C between washes), washed once more in 1 mL cold 10% glycerol, and resuspended in 1 mL cold 10% glycerol. For each electroporation, 50 µL of the resuspended culture was mixed with at least 500 ng of purified, dialyzed recombination template **(Table S1)** in a cold 1 mm cuvette and electroporated using a Gene Pulser Xcell instrument (Bio-Rad) with the following settings: 2,500 V, 25 µF, 600 Ω, 1 mm. Immediately following electroporation, cells were recovered in 950 µL BHIS for 3 h at 37 °C without agitation, then pelleted, resuspended in 100 µL BHI, and plated on selective BHI agar.

Electrocompetent *E. faecium* cells harboring pPM260 were prepared by diluting overnight cultures 1:100 in fresh BHI and cultured at 37 °C with shaking. At OD_600_ = 0.3–0.4, isopropyl β-D-thiogalactopyranoside (IPTG) was added to a final concentration of 1 mM and the cultures were incubated further until OD_600_ = 0.6–0.8. Following incubation, cells were pelleted at 5,000 RCF for 10 min at 4 °C, resuspended in 1 mL ice-cold 10% glycerol, and pelleted again at 7,000 RCF for 8 min at 4 °C. The pellet was resuspended in 500 µL lysozyme solution (10 mM Tris [pH 8.0], 20% sucrose, 10 mM EDTA, 50 mM NaCl, 30 µg/mL lysozyme) and incubated at 37 °C for 20–30 min. Cells were then pelleted (7,000 RCF, 8 min, 4 °C) and washed 3–4 times by resuspension in electroporation solution (0.5 M sucrose, 10% glycerol) before a final resuspension in the same solution. For each electroporation, 100 µL of the resuspended culture was mixed with the recombination template **(Table S1)** in a 2 mm cuvette and electroporated using a Gene Pulser II instrument with Pulse Controller Plus (Bio-Rad) with the following settings: 2,500 V, 25 µF, 400 Ω, 2 mm. Immediately following electroporation, 400 µL SBHI medium (0.5 M sucrose in BHI) was added and cells were recovered for 3 h at 37 °C without shaking before plating on selective BHI agar.

Electrocompetent *F. novicida* U112 cells harboring pFNMB2_7, pFNMB2_10, or pFNMB2_12 were prepared by diluting overnight cultures 1:20 in 60 mL MH broth supplemented with 0.1% L-cysteine, 0.1% D-glucose, 0.1% FCS, and 15 µg/mL kanamycin, then cultured at 37 °C with shaking until OD_600_ = 0.7. Expression of the recombineering cassettes was then induced with anhydrotetracycline (aTc) at a final concentration of 100 ng/mL and the cultures were incubated a further 1 h. All subsequent steps were carried out at room temperature. Cells were pelleted and washed twice in 10 mL wash buffer (0.5 M sucrose, 1 mM EDTA), washed twice more in 10 mL 0.5 M sucrose, and resuspended in 200 µL 0.5 M sucrose. For each electroporation, 50 µL of the resuspended culture was mixed with 1,000 ng of the recombination template **(Table S1)** in a 2 mm cuvette, incubated at room temperature for 10 min, and electroporated using a Gene Pulser Xcell instrument (Bio-Rad) with the following settings: 2,500 V, 25 µF, 400 Ω, 2 mm. Immediately following electroporation, cells were resuspended in 1 mL MH broth supplemented with 0.1% L-cysteine, 0.1% D-glucose, and 0.1% FCS and recovered aerobically for 5 h at 37 °C.

Electrocompetent *E. coli* cells were prepared by diluting overnight cultures of the MG1655 recombineering strain 1:100 in 500 mL LB containing 30 µg/mL chloramphenicol and cultured at 37 °C with shaking at 220 rpm. At OD_600_ = 0.4, L-arabinose was added to a final concentration of 2 mM and the cultures were incubated a further 30 min. All subsequent steps were carried out on ice. Cells were pelleted at 4,000 × g for 15 min at 4 °C, washed twice in 50 mL ice-cold 10% glycerol, and resuspended in 1 mL ice-cold GYT buffer (10% glycerol, 0.125% yeast extract, 0.25% tryptone). For each electroporation, 200 µL of the resuspended culture was mixed with 500 ng of purified, dialyzed dsDNA in a 2 mm cuvette and electroporated using a Gene Pulser Xcell instrument (Bio-Rad) with the following settings: 2,500 V, 25 µF, 200 Ω, 2 mm. Immediately following electroporation, cells were recovered in 1 mL SOC medium for 1.5 h at 37 °C with shaking at 220 rpm and plated on selective LB agar.

#### Structural comparison

The three-dimensional structures of the 11’s Gam-like protein and pPM260’s Gp157 were predicted with AlphaFold 3 using the AlphaFold Server (64). The resulting models were superposed onto the experimentally determined crystal structure of the Gam host-nuclease inhibitor protein from *Desulfovibrio vulgaris* (PDB 2P2U, chain A) (65) using the sequence-independent super algorithm in PyMOL (The PyMOL Molecular Graphics System, open-source v2.5, Schrödinger, LLC), and the aligned structures were rendered as cartoons.

#### Bioinformatic screen

The bioinformatic screen was conducted through three iterative searches. First, the Gam protein of *Staphylococcus* phage 11 (WP_000999048.1) was used as a query in a protein–protein BLAST (blastp 2.11.0+) (66,67) against the NCBI nr database (68), restricted to the *Staphylococcaceae* (NCBI taxonomy ID 90964), returning up to 10,000 target sequences per query; scoring and significance parameters were left at the NCBI defaults. The BLAST search was performed using Biopython (69). The XML result file from the search was used together with a custom Python script to retrieve the putative SSAP/recombinases and SSBs associated with every BLAST hit. Briefly, every BLAST hit was searched in the NCBI Identical Protein Groups (IPG) database to recover the genomes encoding that specific sequence. The first genome associated with each sequence in the IPG database was selected, and the script retrieved the amino acid sequences of the two proteins encoded downstream of each BLAST hit. Additionally, for every three proteins (the BLAST hit and the two proteins encoded downstream), the number of entries associated with each protein in the IPG database was recorded as a proxy for the prevalence of that exact sequence among sequenced genomes. The putative SSAP/recombinases extracted by the script were collapsed to unique amino acid sequences, and any sequences containing internal stop codons were discarded. Seven sequences were then manually selected to represent the range of diversity recovered **(Table S2).** In the second round of the screen, the same method was applied to the seven selected sequences. This time, the Python script extracted the proteins encoded upstream (putative Gams) and downstream (putative SSBs) of each BLAST hit. All the putative Gams retrieved across the seven BLAST searches were collapsed, and six sequences were selected to move forward as queries for round 3 of the screen **(Table S2)**. In this round, the two proteins encoded downstream of the putative Gams were extracted. The putative SSAP/recombinases associated with all six BLAST hits were collapsed, and ten of these were selected. In total, sixteen 3-gene loci from the round 1 and round 3 searches were picked to be tested **(Table S2).** A custom Python script was used to select, for each recombinase, a specific gam-recombinase-SSB locus to clone. The script queried each SSAP/recombinase in the NCBI IPG database and retrieved the protein sequences of the genes upstream and downstream of the SSAP/recombinase for every entry returned. Then every identical Gam-SSAP/Recombinase-SSB combination was collapsed and counted. The most frequent Gam-SSAP/Recombinase-SSB combination for each recombinase was selected (except for WP_000139720.1, for which the two most represented combinations were taken) (Table S2). Each 3-gene locus selected was then used as a query in a nucleotide BLAST search to determine whether any bacterium encoding it was available from ATCC. Where an ATCC strain harbored an identical or near-identical 3-gene locus, that genome was selected. All the 3-gene loci selected were cloned under an IPTG-inducible promoter in vector pPM141 for testing **(Table S1).**

#### Colony PCR

Bacterial colonies were resuspended in 30 µL of lysis buffer (250 mM KCl, 5 mM MgCl_2_, 50 mM Tris-HCl at pH 9.0, and 0.5% Triton X-100) supplemented with 200 ng/µL of lysostaphin (Ambi Products LLC). The samples were incubated for 20 min at 37 °C and then for 10 min at 98 °C in a thermocycler. Following lysis, 0.5 µL of each lysate was used as a template for PCR with TopTaq master mix (Qiagen) and the appropriate primers.

#### Determining gene requirements for recombineering of the SppErf system

*S. aureus* RN4220 and sPM05, each harboring pPM141, pPM260, pAV493, pAV494, or pAV495, were electroporated with 500 ng of PCR product or with 1,000 ng (approximately 38 pmol) of the oligonucleotide PM1370, respectively. Electroporated cells were resuspended in 900 µL of BHIS and recovered for 3 h at 37 °C with shaking at 220 rpm. To quantify total CFU, each sample was serially diluted and plated on BHI agar plates. Additionally, aliquots were plated on BHI plates supplemented with 5 µg/mL chloramphenicol to quantify the number of recombined CFU. The dsDNA substrate used for electroporation was amplified from sPM03 gDNA using Phusion High-Fidelity polymerase (Thermo Scientific) and the primers PM637 and PM638, followed by PCR purification (QIAquick PCR purification kit, Qiagen) and dialysis against water on 0.025 µm nitrocellulose filters (Millipore).

#### Effect of oligonucleotide amount on recombineering efficiency

S. *aureus* sPM05 harboring pPM260 was electroporated with the following amounts of the oligonucleotide PM1370: 250 ng (10 pmol), 500 ng (20 pmol), 750 ng (30 pmol), and 5,000 ng (200 pmol); water was used as a negative control. The samples were recovered in BHIS for 3 h at 37 °C with shaking at 220 rpm. Each sample was serially diluted and plated on BHI agar plates supplemented with 10 µg/mL erythromycin to quantify the total number of CFU and the rest was plated on 5 µg/mL chloramphenicol to quantify the number of recombined CFU. The plates were incubated overnight at 37 °C.

#### Effect of oligonucleotide length on recombineering efficiency

*S. aureus* sPM10 harboring the vector pPM260 was electroporated with 5,000 ng of the following oligonucleotides: PM1585, PM1586, and PM1587; water was used as a negative control. The samples were recovered in BHIS for 3 h at 37 °C with shaking at 220 rpm. Each sample was serially diluted and plated on BHI agar plates supplemented with 10 µg/mL erythromycin to quantify the total number of CFU and the rest was plated on 5 µg/mL chloramphenicol to quantify the number of recombined CFU. The plates were incubated overnight at 37 °C.

#### Generating dsDNA substrates for recombineering

dsDNA substrates were generated by amplifying either plasmid DNA or gDNA using Phusion High-Fidelity polymerase (Thermo Scientific). Plasmid-derived PCR products were purified using a QIAquick gel extraction kit and gDNA-derived PCR products using a QIAquick PCR purification kit (Qiagen). The purified PCR products were then dialyzed against water on 0.025 µm nitrocellulose filters (Millipore).

#### Impact of phosphorothioate-modified oligonucleotides on recombineering efficiency

*S. aureus* sPM10 harboring the vector pPM260 was electroporated with 5,000 ng of oligonucleotide PM1585; water was used as a negative control. PM1585 was used either unmodified or with phosphorothioate linkages at its four 5′-terminal or four 3′-terminal bases. The samples were recovered in BHIS for 3 h at 37 °C with shaking at 220 rpm. Each sample was serially diluted and plated on BHI agar plates supplemented with 10 µg/mL erythromycin to quantify the total number of CFU and the rest was plated on 5 µg/mL chloramphenicol to quantify the number of recombined CFU. The plates were incubated overnight at 37 °C.

#### IPTG induction time experiment

An overnight culture of S. *aureus* RN4220 harboring pPM260 was diluted 1:100 in BHIS supplemented with 10 µg/mL erythromycin in five flasks. The flasks were incubated for a total of 1 h 45 min at 37 °C with shaking at 220 rpm. For each flask, isopropyl β-D-thiogalactopyranoside (IPTG) was added at a final concentration of 1 mM at the following times during the incubation: 45 min, 1 h, 1 h 15 min, and 1 h 30 min. As a control, no IPTG was added to the fifth flask. After 1 h 45 min of incubation, the OD_600_ of each culture was approximately 0.8, and cells were made electrocompetent as described above. A 100 µL aliquot of the induced and non-induced cultures was electroporated with 500 ng of PCR product and then resuspended in 900 µL of BHIS and incubated for 3 h at 37 °C and 220 rpm. To quantify total CFU, each sample was serially diluted and plated on BHI agar plates. Additionally, aliquots were plated on BHI plates supplemented with 5 µg/mL chloramphenicol to quantify the number of recombined CFU. The dsDNA substrate used for electroporation was amplified from sPM03 gDNA using the primers PM637 and PM638.

#### Oligonucleotide strand polarity and MMR evasion

S. *aureus* sPM05 harboring pPM260 was electroporated with 500 ng of each of the following oligonucleotides: PM1367, PM1368, PM1369, and PM1370; water was used as a negative control. The samples were recovered in BHIS for 3 h at 37 °C with shaking at 220 rpm. Each sample was serially diluted and plated on BHI agar plates supplemented with 10 µg/mL erythromycin to quantify the total number of CFU and the rest was plated on 5 µg/mL chloramphenicol to quantify the number of recombined CFU. The plates were incubated overnight at 37 °C.

#### Optimizing the amount of dsDNA for recombineering

gDNA from *S. aureus* sPM03 was PCR amplified with the primers PM637/PM638. Different amounts (250, 500, 1,000, 2,000, or 3,000 ng) of the resulting PCR product were electroporated into *S. aureus* harboring the pPM260 plasmid. Immediately after electroporation, 900 µL of BHIS was added and the samples were recovered for 3 h at 37 °C and 220 rpm. Then, each sample was serially diluted and plated on BHI agar plates supplemented with 10 µg/mL erythromycin to quantify the total number of CFU and an aliquot of each culture was plated on 5 µg/mL chloramphenicol to quantify the number of recombined CFU. The plates were incubated overnight at 37 °C.

#### Phylogenetic analysis of recombineering-cassette proteins

Proteins recovered from the bacteriophage recombineering-cassette screen were re-annotated by re-fetching each protein sequence from the NCBI protein database (E-utilities) (68) and searching it against the PHROG (53) v4 profile database with MMseqs2 (70) (mmseqs search -s 8.5 -e 1e-3, profiles as queries), as implemented in pharokka (55), retaining the best hit per protein by bit score. Restricting to complete cassettes (Gam, recombinase, and single-stranded DNA-binding (SSB) protein all present), a separate tree was built for each cassette position from the proteins assigned to the relevant PHROG groups: the Gam position from PHROG groups 946, 253, and 2205; the recombinase position from groups 2324, 155 (Erf-like), 659 (Sak-like), and 124 and 32104 (Sak4-like); and the SSB position from groups 44 and 378. For each position, sequences were dereplicated to exact-unique sequences and clustered at 90% identity with CD-HIT (71) (v4.8.1) to obtain a representative set. To place the experimentally tested constructs precisely, the exact amino acid sequences of the cloned candidate Gams, recombinases, and SSBs were extracted from their plasmid maps (or, where unannotated, from the corresponding NCBI record) and grafted into the representative set as additional leaves. Each combined set was aligned with MAFFT (72) (v7.526, --auto), trimmed with trimAl (73) (v1.5.1, -gt 0.2), and a maximum-likelihood tree was inferred with IQ-TREE (74) (v3.1.3), selecting the best-fit substitution model with ModelFinder (75) (-m MFP; selected models: Q.PFAM+F+G4 for Gam, LG+F+G4 for recombinase, and Q.INSECT+F+I+R5 for SSB) and assessing branch support with 1,000 ultrafast bootstrap replicates. Trees were midpoint-rooted and visualized as circular phylograms with ggtree (76) (v4.0.5), coloring tips by PHROG group and scaling tip size by the number of genomes represented by each 90%-identity cluster. For the SSB tree, two candidate proteins whose third-cassette gene is not a canonical SSB (pPM256, an HNH endonuclease; pPM263, unclassifiable) were excluded from the final tree.

#### Plasmid construction

All the plasmids used in this study can be found in **Table S1**. The table also includes the cloning strategy for each plasmid and the oligos used to build them. Plasmids were isolated using a QIAprep Spin Miniprep Kit according to the manufacturer’s protocol for *E. coli* and using a modified protocol for *S. aureus* as previously described elsewhere (77).

#### Plasmid curing

To quantify loss of the recombineering plasmid, a single colony of S. aureus sPM03 carrying pPM145v1 was used to inoculate three cultures of BHI supplemented with 5 µg/mL chloramphenicol and containing 0.2, 1.0, or 5.0 mM IPTG, and incubated overnight at 37 °C with shaking at 220 rpm. Each culture was then streaked onto BHI agar supplemented with 5 µg/mL chloramphenicol and incubated overnight at 37 °C. Fifty individual colonies from each IPTG treatment were then patch-plated on BHI agar plates supplemented with either 5 µg/mL chloramphenicol or 10 µg/mL erythromycin. On the basis of these results, all subsequent plasmid curing was performed by growing the relevant strain overnight in BHI supplemented with 0.5 mM IPTG in the absence of antibiotic selection for the recombineering vector.

#### Recombineering in *S. aureus* clinical isolates

Electrocompetent *S. aureus* USA300, M06, MW2, HER1049, and MRSA192 harboring pPM260 were electroporated with 500 ng of a PCR product harboring either a chloramphenicol or spectinomycin (for USA300) resistance cassette flanked with 500 bp of homology to the *sau1hsdR* locus. Following electroporation, the cells were resuspended in 900 µL of BHIS, and then recovered for 3 h at 37 °C with shaking at 220 rpm. The cells were plated on BHI agar plates supplemented with 5 µg/mL chloramphenicol or 125 µg/mL spectinomycin and incubated overnight at 37 °C. The following day, editing was assessed by colony PCR with primers PM637/PM638 with the products resolved on a 1% agarose gel stained with ethidium bromide. The electroporated PCR products were generated by amplification of sPM03 and sPM04 gDNA with primers PM637/PM638.

#### Recombineering with Cre-mediated excision of the antibiotic cassette

Electrocompetent *S. aureus* RN4220 harboring pPM260 or pPM300 (100 µL) was electroporated with 500 ng of PCR product and immediately resuspended in 900 µL of BHIS, then recovered for 3 h at 37 °C with shaking at 220 rpm. The electroporated PCR product was generated by amplification of sPM09 gDNA with primers PM637 and PM638. Following electroporation, each sample was serially diluted and plated on BHI agar plates supplemented with 10 µg/mL erythromycin to quantify the total number of CFU, and an aliquot was plated on BHI agar with 5 µg/mL chloramphenicol to quantify the number of edited CFU. The plates were incubated overnight at 37 °C. To excise the cassette, an edited colony from *S. aureus* RN4220 harboring pPM300 was used to inoculate an overnight culture in BHI supplemented with 5 µg/mL chloramphenicol and 10 µg/mL erythromycin. The overnight culture was diluted 1:100 into two tubes of BHI with 10 µg/mL erythromycin and cultured at 37 °C with shaking at 220 rpm for 1 h 45 min. Anhydrotetracycline (aTc) was then added to one of the tubes at a final concentration of 125 ng/mL to induce *cre* recombinase expression and both tubes were incubated for a further 5 h before streaking on BHI agar plates supplemented with 10 µg/mL erythromycin. The following day, excision was assessed by colony PCR with primers PM651/PM652 on 10 colonies from each culture, with the products resolved on a 1% agarose gel stained with ethidium bromide and confirmed by Sanger sequencing (Genewiz). To confirm the removal of the chloramphenicol cassette and to test for pPM300 curing, a single aTc-treated colony showing excision was used to inoculate three overnight cultures (BHI alone, BHI with 5 µg/mL chloramphenicol, or BHI with 0.5 mM IPTG) incubated at 37 °C with shaking at 220 rpm. To assess IPTG-induced plasmid curing, the untreated and IPTG-treated cultures were then streaked on BHI agar plates. Fifty colonies from each plate were patch-plated on BHI agar with and without 10 µg/mL erythromycin and incubated overnight at 37 °C.

#### Strain construction

*S. aureus* sPM03 was isolated from a single edited colony using pPM145v1 in the 11 recombineering experiment described above. *S. aureus* sPM04, sPM05, sPM09, and sPM10 were generated by recombineering of *S. aureus* RN4220. Briefly, 500 ng of linear DNA was electroporated into *S. aureus* RN4220 harboring the pPM260 plasmid. Following electroporation, an edited colony was isolated on a BHI agar plate supplemented with 125 µg/mL spectinomycin for sPM04, sPM05, and sPM10 or 5 µg/mL chloramphenicol for sPM09. To generate sPM04, sPM05, and sPM10, the electroporated DNA was amplified from the plasmids pPM165, pPM269, and pPM319 using primers PM637 and PM638. For sPM09, a linear DNA fragment was produced using Gibson assembly (78) of three PCR products from the PCR amplification of sPM03 gDNA using the following primer pairs: PM637/PM1462, PM1463/PM1461, and PM638/PM1464.

#### Testing different homology arm lengths

PCR products harboring a chloramphenicol cassette flanked with homology arms of different lengths to the *sau1hsdR* locus were generated by amplifying sPM03 gDNA using the following primer pairs: PM635/PM636 (40 bp), PM1377/PM1378 (60 bp), PM649/PM650 (100 bp), PM651/PM652 (200 bp), PM637/PM638 (500 bp), and PM1372/PM1375 (2 kb). Then, 2.7 pmol of each PCR product was electroporated into 100 µL of electrocompetent *S. aureus* RN4220 carrying the pPM260 vector. The cells were then resuspended in 900 µL of BHIS and incubated for 3 h at 37 °C and 220 rpm. Each sample was serially diluted and plated on BHI agar plates supplemented with 10 µg/mL erythromycin to quantify the total number of CFU. Additionally, aliquots were plated on BHI plates supplemented with 5 µg/mL chloramphenicol to quantify the number of recombined CFU.

#### Testing the recombineering cassettes from the bioinformatic screen

*S. aureus* carrying an empty plasmid control (pPM141), the SapSak4 system or one of the selected 3-gene recombineering cassettes was electroporated with 500 ng of PCR product amplified from sPM03 gDNA with PM637/PM638. Immediately after electroporation, 900 µL of BHIS was added and the samples were recovered for 3 h at 37 °C and 220 rpm. Then, 50 µL of each sample was serially diluted and plated on BHI agar plates supplemented with 10 µg/mL erythromycin to quantify the total number of CFU and the rest was plated on 5 µg/mL chloramphenicol to quantify the number of recombined CFU. The plates were incubated overnight at 37 °C.

#### Prevalence of Gam–SSAP–SSB cassettes in staphylococcal phages

To estimate the genome-wide prevalence of the recombineering cassette independently of the synteny-seeded screen, all complete staphylococcal phage genomes were retrieved from the INPHARED database (79) (9 August 2025 release; n = 1,018 records with Host = *Staphylococcus*). Nucleotide sequences were downloaded from NCBI by accession (Biopython v1.88 Entrez). Genomes were annotated with Pharokka v1.10.1 (55) in metagenome mode (-m; genes predicted with Prodigal-gv via pyrodigal-gv v0.3.2) against the PHROG-based Pharokka database v1.8.0 (53), yielding 116,366 CDS. Each CDS was then structurally re-annotated with Phold v1.2.6 (E-value ≤ 1 × 10⁻³, sensitivity 9.5). Recombineering cassettes were identified from the Phold per-CDS annotations by gene synteny: a single-strand annealing protein (SSAP) was called a cassette recombinase when immediately flanked (±1 CDS) by a Gam/Gp157 end-protection protein, and the cassette was scored as complete when a single-stranded DNA-binding protein (SSB) occupied the CDS on the opposite side of the SSAP, strand-aware in the Gam-SSAP-SSB orientation. SSAP families (Erf, Sak4, Sak, RecT, Gp2.5) were assigned from Phold product annotations. Prevalence was reported as the fraction of the 1,018 genomes carrying each feature.

#### SapSak4 recombineering experiment

*S. aureus* RN4220 (80) harboring one of five plasmids (pE194, pPM145v1, pPM145v4, pPM145v6, or pPM145v8) was electroporated using 500 ng of a PCR product harboring a chloramphenicol acetyltransferase cassette flanked on each side by 500 bp of homology to the non-functional *sau1hsdR* gene of RN4220 (23,81). The electroporated DNA was generated by PCR of the pPM146 vector with the primer pair PM637/PM638. After electroporation, 800 µL of BHIS was added to the cultures, which were recovered for 2 h at 37 °C with shaking at 220 rpm. Following recovery, 850 µL of culture was pelleted, resuspended in 100 µL of BHI and plated on BHI agar plates supplemented with 5 µg/mL chloramphenicol and 10 µg/mL erythromycin to quantify the number of edited cells. The leftover 50 µL of culture was serially diluted and plated on BHI agar plates supplemented with 10 µg/mL erythromycin to quantify the total number of colony-forming units (CFU). The plates were then incubated overnight at 37 °C and the number of colonies was quantified the next day. Following quantification, eight colonies (one only, for pPM145v4, which yielded a single colony) from the BHI agar plates supplemented with 5 µg/mL chloramphenicol and 10 µg/mL erythromycin were streaked on fresh plates and incubated overnight at 37 °C. The next day, a colony from each plate was screened by colony PCR for editing of the *sau1hsdR* locus using the primers PM31/PM646. Colonies of *S. aureus* RN4220 were used as controls.

